# Topological analysis of TMEM180, a newly identified membrane protein that is highly expressed in colorectal cancer cells

**DOI:** 10.1101/789750

**Authors:** Takahiro Anzai, Yasuhiro Matsumura

## Abstract

New target molecules for diagnosis of and drug development for colorectal cancer (CRC) are always in great demand. Previously, we identified a new colorectal cancer–specific protein, TMEM180, and successfully developed an anti-TMEM180 monoclonal antibody (mAb) for the diagnosis and treatment of CRC. Although TMEM180 is categorized as a member of the cation symporter family and multi-pass membrane protein, little is known about its function. In this study, we examined topology of this membrane protein and analyzed its function. Using a homology model of human TMEM180, we experimentally determined that the protein has 12 transmembrane domains, and that its N-terminal and C-termini are exposed extracellularly. Moreover, we found that the putative cation-binding site of TMEM180 is conserved among orthologs, and that its position is similar to that of melibiose transporter MelB. These results suggest that TMEM180 acts as a cation symporter. Our topological analysis based on the homology model provides insight into functional and structural roles of TMEM180 that may help to elucidate the pathology of CRC.

**Highlights:**

- A homology model of human TMEM180 was generated by secondary structure prediction.
- Putative cation-binding residues are conserved in TMEM180 orthologs.
- Both the N-terminus and C-terminus of TMEM180 are extracellularly exposed.
- TMEM180 is a 12-transmembrane protein.
- TMEM180 could act as a cation symporter.

## 1. Introduction

Colorectal cancer (CRC) is one of the most common cancers, and has high mortality and incidence rates around the world [1]. Consequently, there is a considerable incentive to identify new target molecules for diagnosis of and drug development for this disease. We previously identified a new CRC-specific protein, TMEM180, and developed the anti-TMEM180 monoclonal antibody (mAb) for use in diagnosing and treating CRC [2]. TMEM180 is also called MFSD13A (major facilitator superfamily domain–containing 13A), but the recommended protein name in the UniProtKB database (entry Q14CX5) remains transmembrane protein 180 [3]. The TMEM180 gene encodes a transmembrane protein that belongs to the glycoside–pentoside–hexuronide (GPH):cation symporter family (Transporter Classification Database, http://www.tcdb.org/) [4]. Members of this family catalyze symport of a sugar molecule with a monovalent cation (H^+^ or Na^+^) (http://www.tcdb.org/search/result.php?tc=2.A.2) [4]. TMEM180 is categorized as a member of the cation symporter family and a multi-pass membrane protein, but little information is available regarding its substrate and topology (http://www.tcdb.org/search/result.php?tc=2.A.2.3.10) [4].

In order to elucidate the functional and structural role of TMEM180, we decided to analyze the topology of TMEM180. Here, we present our *in silico* analysis of TMEM180 and a homology model based on secondary structure prediction. We also present a reliable experimental topological model of TMEM180. We show that the both N-terminus and C-terminus of TMEM180 are extracellularly exposed. Immunofluorescence with or without cell membrane permeabilization was performed to confirm the topological model of TMEM180. We found that the putative cation-binding site in TMEM180 is highly conserved among TMEM180 orthologs. Our topological analysis based on the homology model provides insight into the functional and structural roles of TMEM180, which may help elucidate the pathology of CRC.

## 2. Materials and Methods

### 2.1. Cells and Cell culture

Human embryonic kidney (HEK) 293T cells were purchased from American Type Culture Collection. HEK293T cells were cultured in DMEM (Wako) supplemented with 10% FBS (Thermo Fisher Scientific) and 1% penicillin–streptomycin–amphotericin B suspension (Wako) at 37°C under a 5% CO_2_ atmosphere.

### 2.2. Plasmid construction

The human *TMEM180* gene was PCR amplified from plasmid pCMV-TMEM180 (OriGene) and fused with an *Eco*R1 restriction site in the 5’ region and a *Bam*H1 restriction site in the 3’ region. The PCR products were cloned into pcDNA3.3-TOPO (Thermo Fisher Scientific), and the resultant plasmid was designated pcDNA3.3-hTMEM180. The human *TMEM180* genes with an N-terminal 3*FLAG-tag, C-terminal FLAG-tag, or N-terminal HA-tag were amplified and fused by overlap extension PCR and cloned into the *Eco*R1/*Bam*H1 restriction sites of pcDNA3.3-hTMEM180 using In-Fusion cloning technology (Clontech). FLAG-tag insertion constructs were generated by inverse PCR or overlap extension PCR using the In-Fusion cloning technology. All of the plasmid constructs used in this study were verified by DNA sequencing. Primers used for plasmid construction in this study are described in Supplementary Table1.

### 2.3. Immunofluorescence

HEK293T cells were transfected for 36–48 h with individual TMEM180 plasmids using Lipofectamine LTX (Thermo Fisher Scientific). After transfection, cells were washed with PBS and fixed with 4% PFA/PBS (Wako) for 15 min. After the treatment of the cell membrane with PBS or 0.1% Triton X-100/PBS for 10 min, cells were blocked with 5% skim milk/PBS and stained with primary antibody: humanized anti-TMEM180 mAb (2 µg/mL, clone 669, RIN Institute), anti-FLAG mouse mAb (1:200, clone M2, Sigma-Aldrich), anti-FLAG mouse mAb (1:200, clone M5, Sigma-Aldrich), or anti-HA rabbit pAb (1:200, MBL). The cells were then incubated with the appropriate secondary antibody: Alexa Fluor 488–conjugated anti–human IgG goat mAb (1:200, Thermo Fisher Scientific), Alexa Fluor 647–conjugated anti–mouse IgG goat mAb (1:200, Thermo Fisher Scientific), or Alexa Fluor 488–conjugated anti–rabbit IgG goat mAb (1:200, Thermo Fisher Scientific). The cells were also incubated with DAPI (2 µg/mL, Thermo Fisher Scientific) to stain nuclear DNA. Images were acquired on a BZ-X710 fluorescence microscope (Keyence) and analyzed using ImageJ version 1.52a (NIH).

### 2.4. *in silico* analysis

The full-length amino acid sequence of human TMEM180 was retrieved from UniProtKB entry number Q14CX5 (1–517 amino acids). The online programs MEMSAT-SVM [5], PRODIV-TMHMM [6], TMHMM [7], MemBrain [8], Phobius [9], Philius [10], TMpred [11], DAS-TMfilter [12], HMMTOP [13], CCTOP [14], SOSUI [15], Scampi [16], Scampi-MSA [16], and PRO-TMHMM [17], with data from UniProt [3], were used for prediction of transmembrane regions of human TMEM180 using the full-length amino acid sequence.

### 2.5. Homology modeling

The full-length amino acid sequence of human TMEM180 was retrieved as described in section 2.4. To predict the secondary structure of human TMEM180, we used two different servers, DELTA-FORTE [18] and HHPred [19]. MEDELLER [20] was used to make a homology model of human TMEM180 using the melibiose transporter MelB from *Salmonella typhimurium* (stMelB) (PDBID 4M64) chain A as the template structure. Target–template alignment from DELTA-FORTE (Fig. S1) [18] was used as input for the MEDELLER workspace. Multiple sequence alignment of TMEM180 from human, bovine, mouse, chicken, frog, and zebrafish corresponding to UniProtKB entry numbers Q14CX5, Q58CT4, Q6PDE8, Q5ZKJ5, B1H1F7, and Q5PRC2 was performed using Clustal Omega (https://www.ebi.ac.uk/Tools/msa/clustalo/) [21]. Evolutionarily conserved amino acid residues of TMEM180 were identified using the ConSurf Server (http://consurf.tau.ac.il/) [22]. All structures in this study were visualized using PyMol version 1.3 [23].

## 3. Results

### 3.1. *In silico* analysis of human TMEM180

To determine the topology of TMEM180 in the plasma membrane, we first performed an *in silico* analysis to investigate the transmembrane (TM) regions of human TMEM180 (UniProtKB entry number Q14CX5) using 14 different prediction programs: MEMSAT-SVM [5], PRODIV-TMHMM [6], TMHMM [7], MemBrain [8], Phobius [9], Philius [10], TMpred [11], DAS-TMfilter [12], HMMTOP [13], CCTOP [14], SOSUI [15], Scampi [16], Scampi-MSA [16], and PRO-TMHMM [17]. Data were obtained from UniProt [3]. According to this analysis, the number of TM regions in human TMEM180 was estimated to be between 8 and 12 (Fig. 1A). TM1, TM9, and TM12 were predicted by all programs, and TM3, TM5, and TM6 were predicted by all programs except one (Fig. 1A,B). Of the 12 TMs, 6 were predicted with high accuracy, whereas the other half were difficult to predict. There was no regularity of the positions of the TMs that were predicted with (Fig. 1B). Consequently, we decided to use secondary structure prediction and construct a homology model to investigate the topology of human TMEM180.

**Fig. 1.**
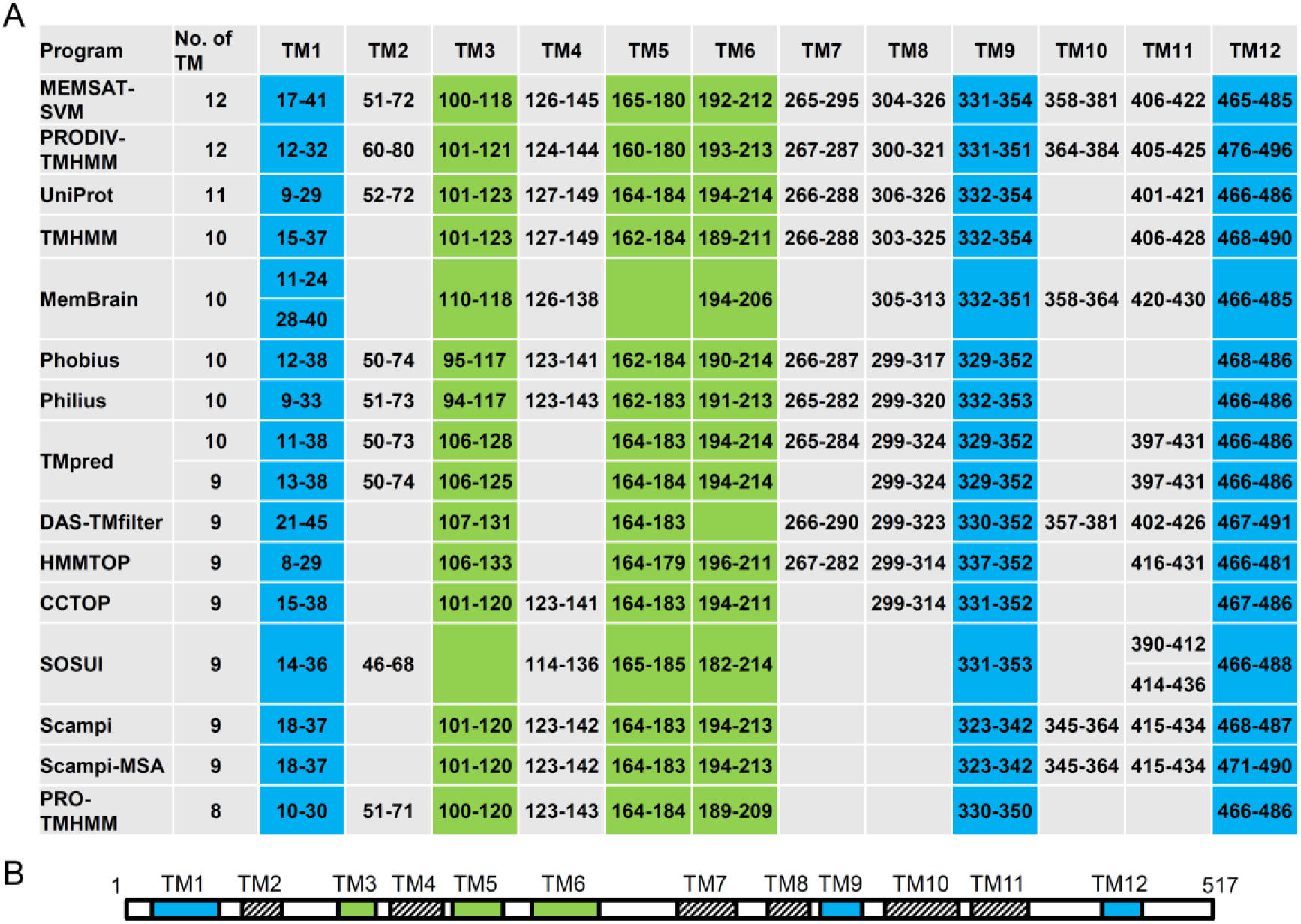
Prediction of transmembrane regions of human TMEM180. A, Transmembrane regions of human TMEM180 were predicted using 14 prediction programs and one database. The numbers indicate TM regions predicted by each program. The regions highlighted in blue were predicted by all prediction programs, and those in green were predicted by all programs except one. B, Schematic models of transmembrane regions for human TMEM180 using most predicted TM number. The regions were highlighted using the color scheme described in Fig. 1A.

### 3.2. Homology modeling of human TMEM180

To address the function of TMEM180 based on structural information, we constructed a homology model of TMEM180. To this end, the primary sequence of human TMEM180 was submitted to the protein BLAST server at NCBI (https://blast.ncbi.nlm.nih.gov/Blast.cgi?PAGE=Proteins), using the algorithms BLASTP, PSI-BLAST, and PHI-BLAST for the homology search. Unfortunately, we could not find any homologous sequences, except for TMEM180 orthologs from other species. Next, we performed secondary structure prediction using DELTA-FORTE [18] and HHpred [19] independently. Both algorithms returned the melibiose transporter MelB from *Salmonella typhimurium* (stMelB) (PDBID 4M64) [24] (Fig. 2A, right) chain A as the top-scoring hit. We then used the on-line program MEDELLER [20], which is specialized for homology modeling of membrane proteins, and adopted stMelB chain A as the template structure. We obtained a homology model of human TMEM180 containing 12 TM (Fig. 2A, left). Although human TMEM180 has only 19% sequence identity with stMelB, comparison of the crystal structure of stMelB with the homology model of human TMEM180 revealed that five putative cation-binding residues (Asn63, Asn66, Asp67, Thr139, and Asp142) in the homology model were located in positions similar to those of five cation-binding residues of stMelB (Asp55, Asn58, Asp59, Thr121, and Asp124) [24,25] (Fig. 2A,B). Multiple sequence alignment of TMEM180 from human, bovine, mouse, chicken, frog, and zebrafish revealed that these putative cation-binding residues are conserved across all species (Fig. 2B). In addition, we used the ConSurf Server [22] to identify evolutionarily conserved amino acid residues of TMEM180 from human, bovine, mouse, chicken, frog, and zebrafish. Mapping of the results onto the homology model of human TMEM180 revealed that the core helix regions are highly conserved, whereas the long loops are not (Fig. 2C). These results indicate that TMEM180 may have 12 TM regions and may act as a cation symporter. We then examined to verify this homology model experimentally.

**Fig. 2.**
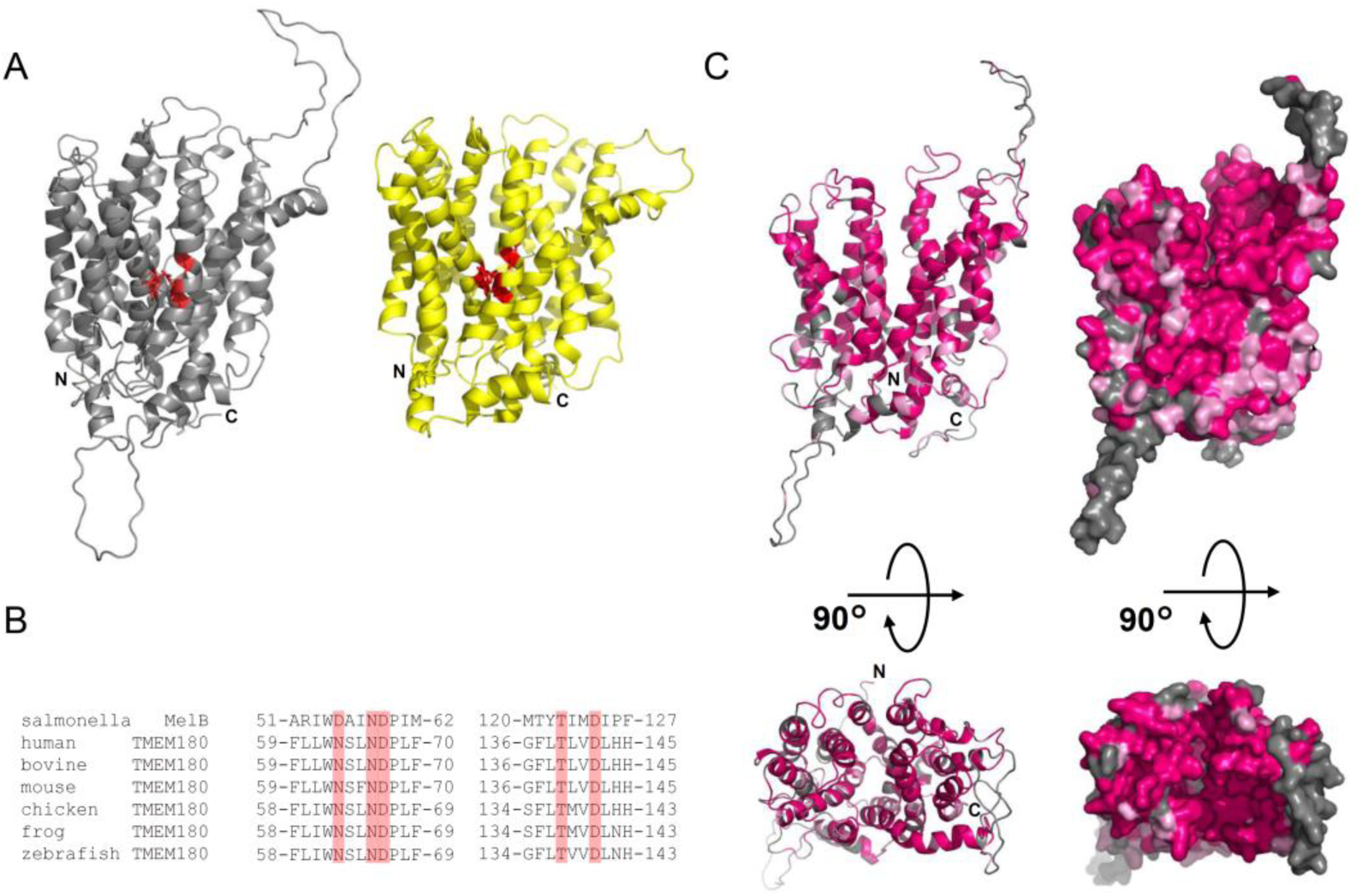
Homology modeling of human TMEM180 and amino acid sequence analysis based on the homology model. A, Homology model of human TMEM180 (gray, left) and the crystal structure of stMelB (yellow, right, PDB ID:4M64). Putative cation-binding sites related to Fig. 2B are colored in red. N, N-terminal; C, C-terminal. B, Multiple sequence alignment based on secondary structure near the cation-binding site of stMelB and TMEM180 from human, bovine, mouse, chicken, frog, and zebrafish. Conserved putative cation-binding residues are highlighted in red C, Amino acid identity (magenta) and similarity (pink) of TMEM180 orthologs shown in (B) are mapped onto the homology model. A ribbon model (left) and surface model (right) are shown. N, N-terminal; C, C-terminal.

### 3.3. N-terminal and C-terminal topology of human TMEM180

First, we decided to determine the N- and C-terminal topology of human TMEM180. For this purpose, FLAG-tags were introduced at the N- or C-terminus of TMEM180 (Fig. 3B,D), and HEK293T cells were transiently transfected with the FLAG-tagged proteins. HEK293T cells transfected with N-terminally 3xFLAG-tagged TMEM180 were stained positively with both anti-TMEM180 mAb and anti-FLAG mAb after permeabilization (Fig. 3A,B upper panel). Non-permeabilized cells were also stained positively by both antibodies (Fig. 3A,B lower panel). Similarly, cells transfected with the C-terminally 3xFLAG-tagged construct were stained by both antibodies whether the cells were permeabilized (Fig. 3C,D upper panel) or not permeabilized (Fig. 3C,D lower panel). These results indicate that both the N- and C-terminus of TMEM180 are extracellularly exposed. Moreover, TMEM180 must have an even number of transmembrane segments in the plasma membrane.

**Fig. 3.**
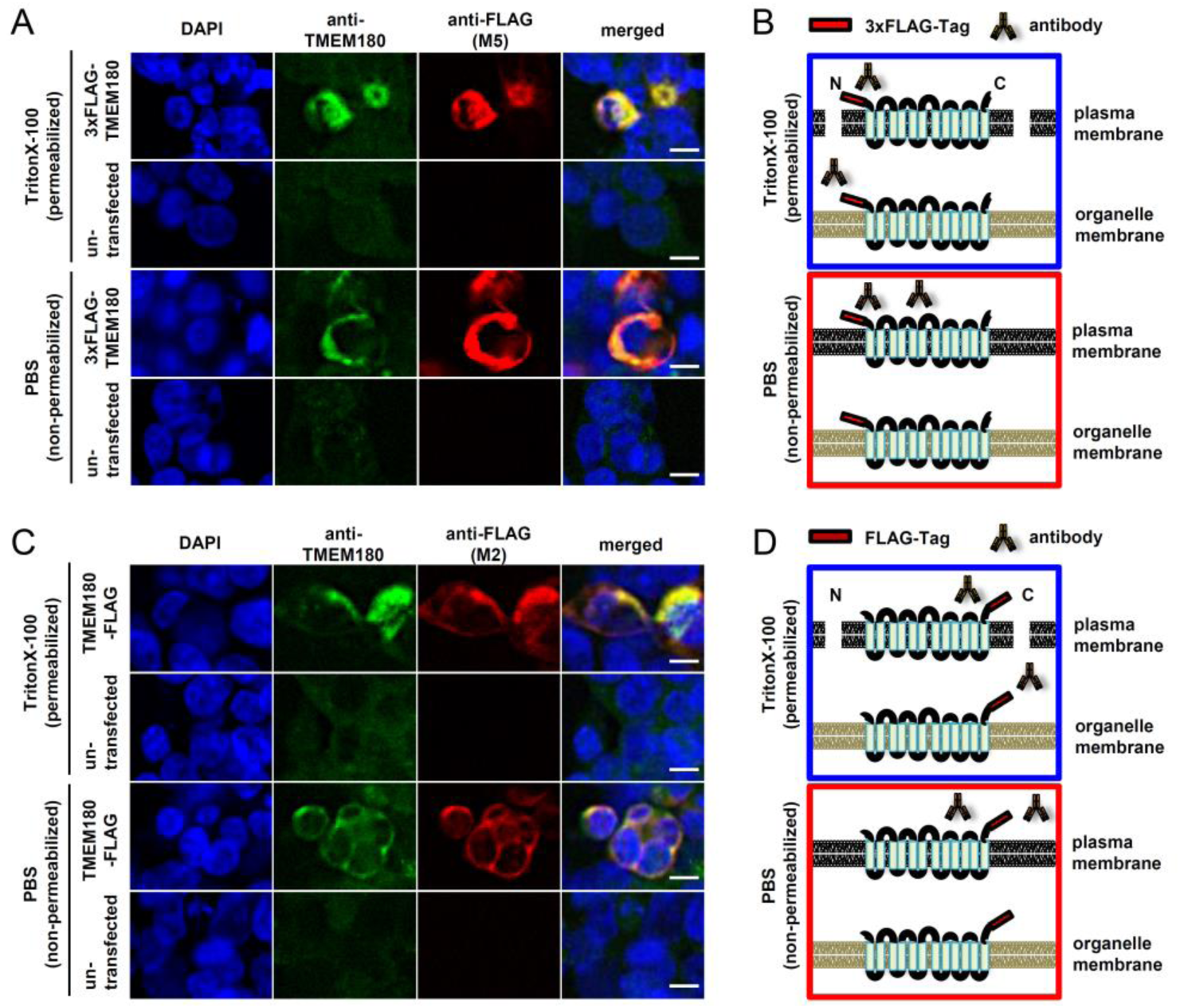
Both the N- and C-termini of TMEM180 are extracellularly exposed. A and C, Immunostaining of HEK293T cells transiently transfected with 3*FLAG-TMEM180 (A) or TMEM180-FLAG (C), either permeabilized (Triton X-100) or not (PBS). HEK293T cells transfected without plasmid were used as negative controls. DAPI (blue), TMEM180 (green), FLAG-tag (red), and merged images are shown. Scale bar = 10 µm. B and D, Schematic diagram of A (B) and C (D) are shown. Non-permeabilized (red boxes) and permeabilized (blue boxes) are shown. Images for each construct were collected under the same conditions. Data are representative of multiple independent experiments.

### 3.4. The entire topology of human TMEM180

To elucidate the topology of TMEM180 in more detail, we constructed TMEM180 genes with an N-terminal HA-tag in the FLAG-tag inserted in loops predicted based on the homology model (Fig. 4A). We chose to use the HA-Tag to confirm expression rather than the anti-TMEM180 mAb that we developed previously [2] because the location of the epitope for this mAb remains unknown. HEK293T cells transfected with a TMEM180 gene with FLAG inserted into five predicted loops (79FLAG, 153FLAG, 236FLAG, 329FLAG, and 398FLAG) were stained by both anti-HA pAb and anti-FLAG mAb regardless of permeabilization (Fig. 4B). On the other hand, HEK293T cells transfected with a TMEM180 gene with FLAG inserted into the other six predicted loops (46FLAG, 123FLAG, 188FLAG, 298FLAG, 357FLAG, and 445FLAG) were stained with both anti-HA pAb and anti-FLAG mAb only in permeabilized HEK293T cells (Fig. 4B). These results indicate that TMEM180 has 12 TMs, and that the topology of our homology model is accurate.

**Fig. 4.**
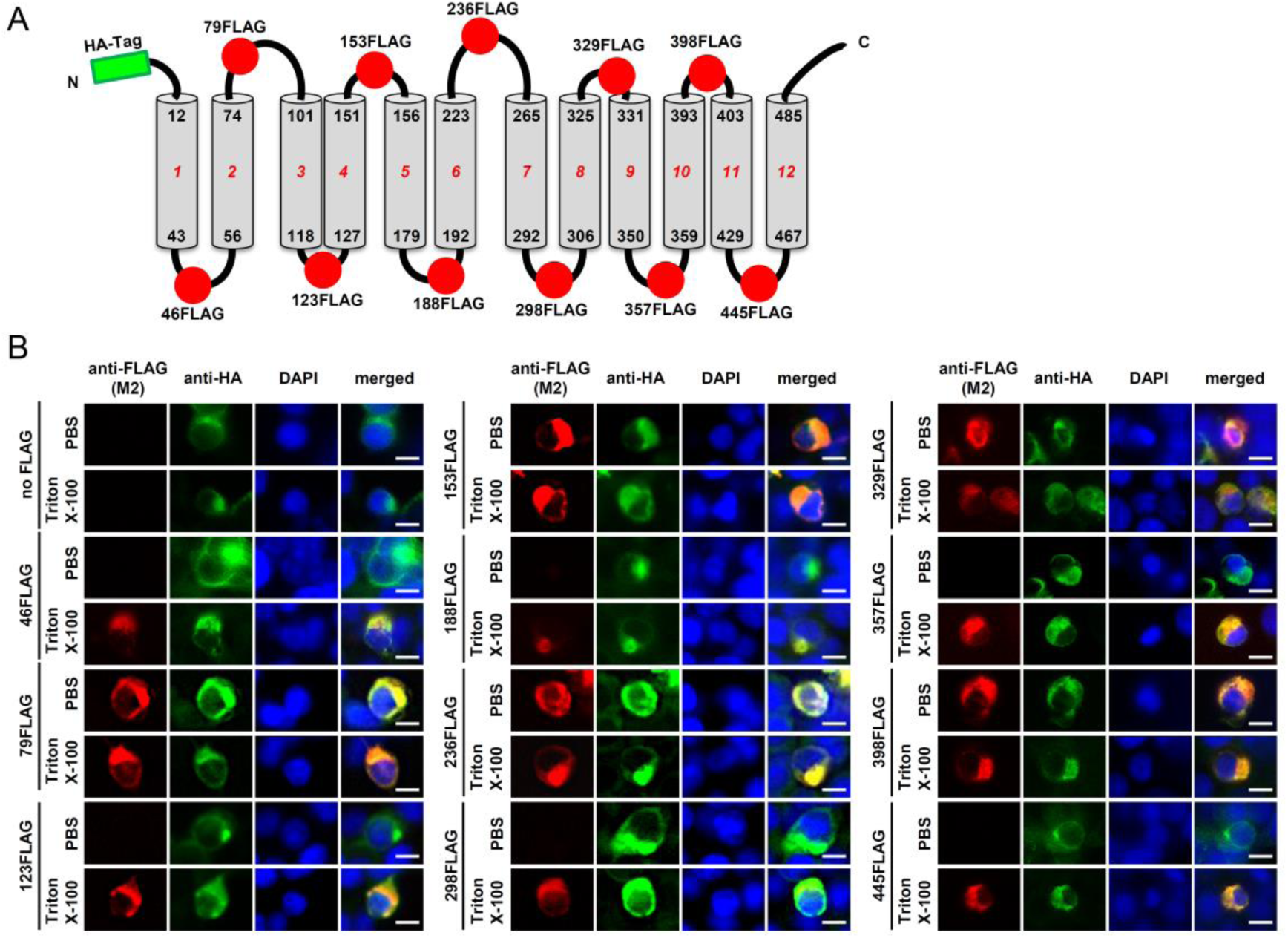
TMEM180 topology mapping using FLAG insert constructs. A, Schematic models of human TMEM180 based on our homology model. Green rectangle represents HA-Tag introduced at the N-terminus, and red circles represent positions of inserted FLAG-tags. N, N-terminal; C, C-terminal. Numbers represent transmembrane residues in the homology model (black) and TM regions (red). B, Immunostaining of HEK293T cells transiently transfected with N-terminally HA-tagged TMEM180 containing FLAG-tag insertions, permeabilized (Triton X-100) or not (PBS). DAPI (blue), HA-tag (green), FLAG-tag (red), and merged images are shown. Scale bar = 10 µm. Data are representative of multiple independent experiments.

## 4. Discussion

The normal counterpart to cancer commonly contains not only normal colonocytes but also fibroblasts, white blood cells, and smooth muscle cells even when laser micro-dissection is used to isolate the normal colonocytes. Therefore, it is difficult to obtain truly cancer-specific molecules under such conditions, even when comprehensive expression analysis is performed. Previously, in our laboratory, we subjected 100% pure normal colonocytes obtained from lavage solution after colonoscopy to comprehensive expression analysis along with colorectal cancer cell lines [26]. Following this analysis, we performed RT-PCR and *in-situ* hybridization; as a result, we identified TMEM180, a CRC-specific multi-pass membrane protein of unknown function [2]. In this study, we analyzed protein structure, including topology, to elucidate the molecular function of TMEM180.

To this end, we constructed a homology model of human TMEM180 (Fig. 2A) using the melibiose transporter MelB which was considered as a structural template. Like TMEM180, MelB belongs to the glycoside–pentoside–hexuronide (GPH):cation symporter family (Transporter Classification Database, http://www.tcdb.org/) [4]. MelB catalyzes symport of melibiose with a monovalent cation. The residues of stMelB involved in cation binding include Asp55, Asn58, Asp59, and Thr121 (Fig. S1, red), whereas those involved in galactoside binding include Asp19, Trp128, Arg149, and Trp342 (Fig. S1, blue); Residues Tyr120, Asp124, and Lys377 (Fig. S1, purple) may be shared by both binding sites [24,25]. In human TMEM180, cation binding residues (Asn63, Asn66, Asp67, and Thr139) and residues shared by the cation and sugar bindings sites (Asp142 and Lys412, but not Leu138) are located in similar positions (Fig. S1, red and purple) and are highly conserved in TMEM180 orthologs [3]. On the other hand, the galactoside-binding residues (Fig. S1, blue) are not conserved at all in TMEM180. Based on our topological analysis, TMEM180 may act as a cation symporter, but does not transport galactosides. Further functional analysis is necessary to determine the substrate of TMEM180.

Previously, we reported that TMEM180 is upregulated under low-oxygen conditions, and that TMEM180 may play an important role in the uptake or metabolism of glutamine and arginine in cancer cell proliferation [2]. We also showed that expression of TMEM180 is not essential for mouse development, as *Tmem180*-knockout mice do not exhibit embryonic, neonatal, or postnatal lethality [2]. During the preparation of this manuscript, a group from Huazhong University of Science and Technology reported that SNP rs2001389 is significantly associated with pancreatic cancer risk, weakens the binding activity of the transcriptional repressor CCCTC-binding factor (CTCF), and is associated with the lower expression of *MFSD13A/TMEM180*. In addition, they showed that siRNA-mediated knockdown of *MFSD13A/TMEM180* promotes proliferation of pancreatic cancer cell lines [27]. On the other hand, we found that shRNA-mediated knockdown of *TMEM180* suppressed proliferation of CRC cell lines (data not shown). Thus, the relationship between expression of TMEM180 and cancer proliferation remains controversial.

This is the first report of the topological structure of TMEM180. We performed *in silico* analysis (Fig. 1A,B) and constructed a homology model of human TMEM180 based on secondary structure prediction (Fig. 2A). We then experimentally confirmed that TMEM180 has 12 TM domains (Fig. 4B) and showed that its N- and C-termini are extracellularly exposed (Fig. 3A,C). Moreover, we found that the putative cation-binding site of TMEM180 is conserved among orthologs, and that the positions of the important residues are similar to those in the melibiose transporter MelB (Fig. 2A,B). These results suggest that TMEM180 acts as a cation symporter. We expect that our model will be useful for analysis of biological function of TMEM180, as well as in determination of its high-resolution x-ray crystal structure or cryo-electron microscopic structure.

## Abbreviations

TMEM180: transmembrane protein 180;
MFSD13A: major facilitator superfamily domain– containing 13A;
mAb: monoclonal antibody;
pAb: polyclonal antibody;
CRC: colorectal cancer;
siRNA: small interfering RNA;
shRNA: small hairpin RNA;

## Acknowledgements

The authors thank members of the Matsumura laboratory for helpful discussion, especially Dr. S. Saijou and Dr. S. Hanaoka for valuable comments, and M. Nakayama and M. Shimada for secretarial support. This work was financially supported in part by JSPS KAKENHI (Grant Numbers JP19K16730 to T.A.) from the Ministry of Education, Culture, Sports, Science and Technology of Japan and the National Cancer Center Research and Development Fund (29-A-9 to Y.M.) from the National Cancer Center, Japan.

## Author Contributions

T.A. and Y.M. conceived the study. T.A. designed and performed all the experiments. T.A. and Y.M. discussed and wrote the manuscript.

---

**Fig. S1.**
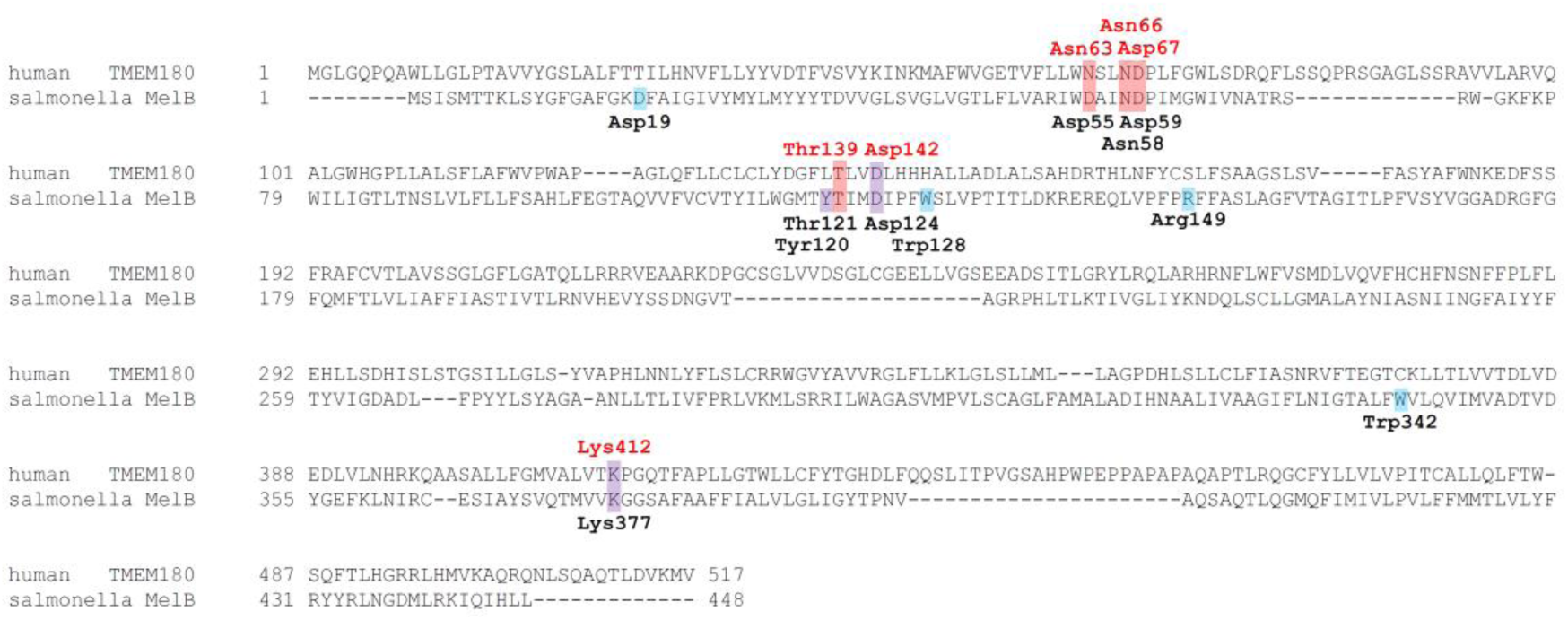
Sequence comparison between human TMEM180 and stMelB based on secondary structure prediction. Sequence alignment of human TMEM180 and stMelB (a.a. 449–487 are missing in the crystal structure) [24] based on secondary structure prediction using DELTA-FORTE [18]. Red represents cation-binding residues, blue represents galactoside-binding residues, and purple represents shared residues. Amino acid numbers from human TMEM180 and stMelB are shown in red and black, respectively.

**Supplemental Table 1.**
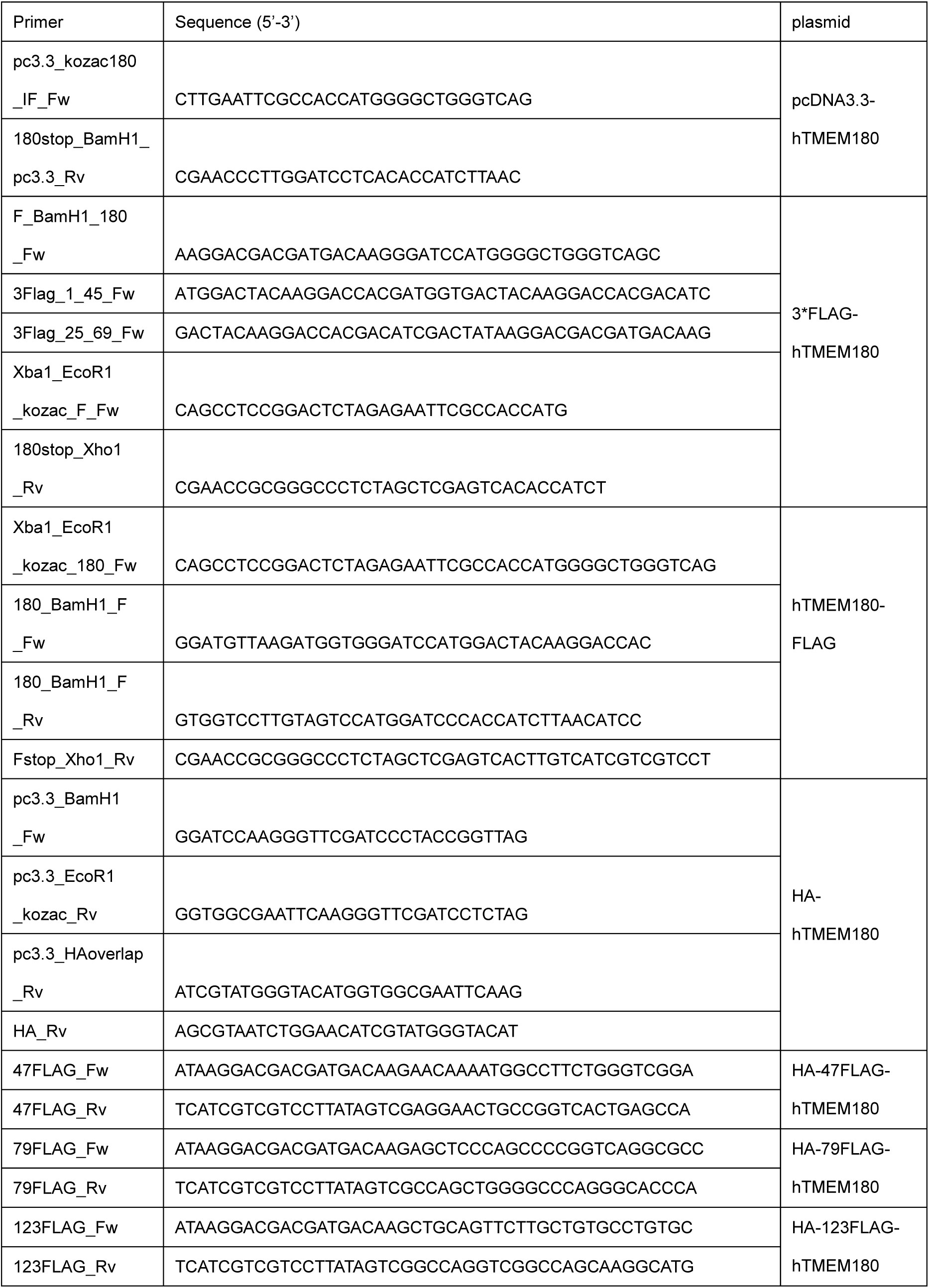
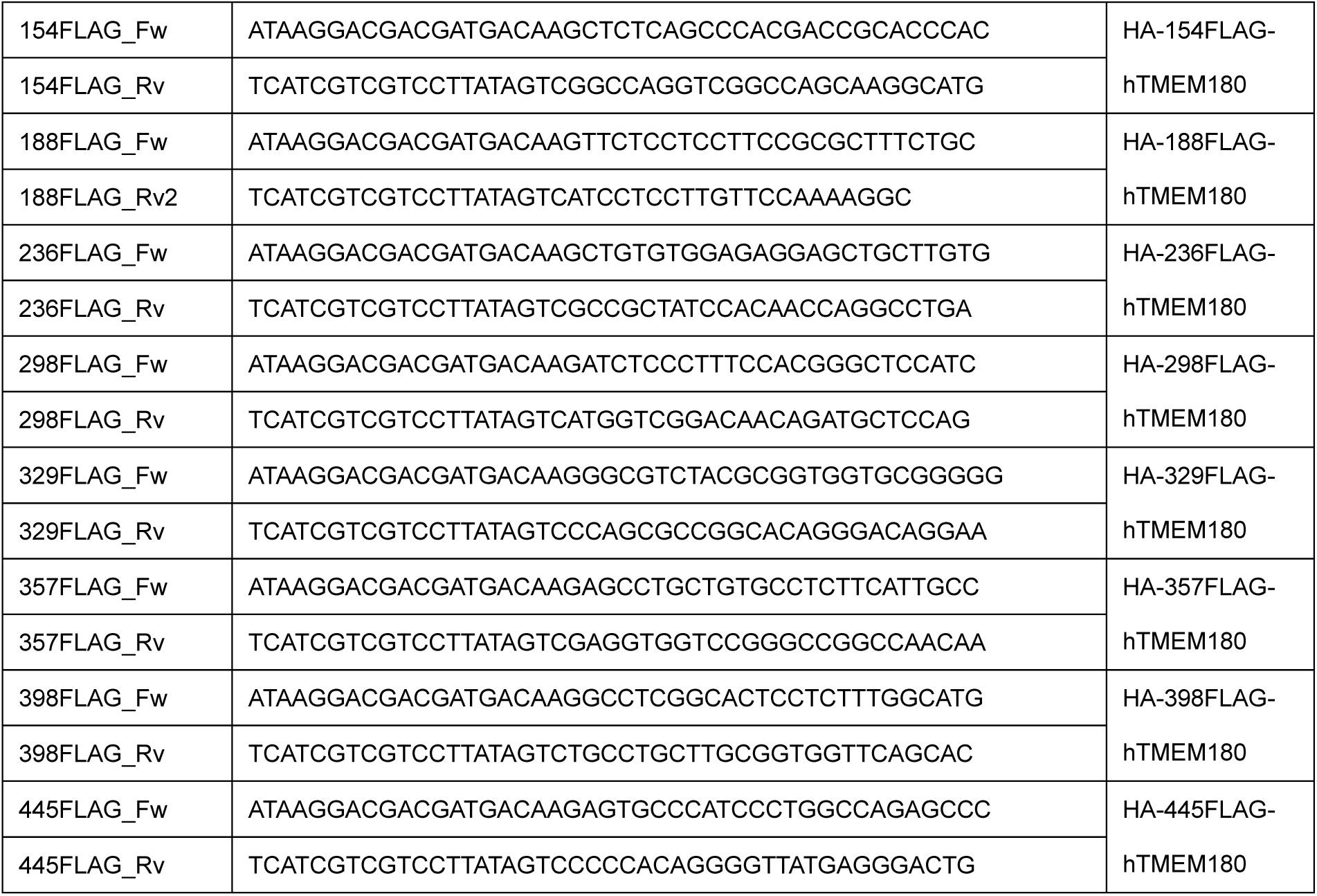
List of primers used in this study

